# sRNAMap: a lightweight, browser-based web application for small RNA mapping, analysis and visualisation

**DOI:** 10.64898/2026.09.22.753470

**Authors:** Melinda Reuter, Lachlan De Hayr, Christopher Chong Wei Ang, Wessel Willemsen, Grant A. Kay, Esther Schnettler, Alain Kohl, Rhys H. Parry

## Abstract

Small RNA (sRNA) sequencing is widely used to study siRNA, miRNA and piRNA pathways and to profile sRNAs derived from viruses, transposons and host transcripts. However, many sRNA workflows rely on command-line mapping and separate software for downstream analysis and visualisation. We present sRNAMap, an HTML5/JavaScript application for mapping and analysing sRNA sequencing data. sRNAMap accepts either raw FASTA/FASTQ reads or pre-aligned BAM files. For raw reads, sRNAMap uses a k-mer–seeded alignment algorithm that runs directly in the browser using parallel Web Workers where supported. It generates strand-specific per-base coverage, 5′- and 3′-end profiles, read-length distributions and nucleotide-bias summaries, with optional library-size normalisation. It also quantifies characteristic read-overlap signatures, including 5′-to-5′ overlaps associated with ping-pong–like processing. Additional modules quantify phased sRNA production, non-templated 3′ and 5′ additions, and sequence diversity. Analyses can be saved and restored as JSON or exported as self-contained HTML reports containing figures and analysis metadata. Benchmarking against Bowtie 2 under approximately matched alignment settings using a representative virus-infected arthropod small-RNA dataset showed highly concordant mapping profiles. Per-position depth showed an R^2^ of approximately 0.998, while 5′- and 3′-end counts showed R^2^ values of approximately 0.97. sRNAMap is available at https://github.com/rhparry/sRNAMap under the MIT licence.

**Impact Statement:** Small RNA sequencing is widely used to investigate antiviral RNAi and other small RNA pathways during virus infection, but analysis often requires command-line mapping followed by multiple separate software packages, creating a barrier to routine exploration and visualisation. We developed sRNAMap to provide an accessible, self-contained alternative that performs small RNA mapping, analysis and visualisation entirely within a web browser. Raw FASTA/FASTQ reads can be mapped directly to user-defined reference sequences, while pre-aligned BAM files can also be imported for downstream analysis. sRNAMap integrates strand-specific coverage, read-length and nucleotide-bias profiling with analyses of read-overlap signatures, phasing, non-templated additions, sequence diversity and nucleotide context. Because all computation occurs locally, sequencing data do not need to be uploaded to an external server. Benchmarking against Bowtie 2 demonstrated highly concordant genome-wide coverage and read-end profiles. sRNAMap therefore provides a lightweight and portable tool for researchers studying virus-derived and other small RNA populations, while lowering the computational barrier to detailed and reproducible small RNA analysis.

## Introduction

Small RNAs (sRNAs) mediate gene regulation and genome defence through several pathways, including small interfering RNA (siRNA), microRNA (miRNA) and PIWI-interacting RNA (piRNA) pathways [1, 2]. Early studies showed that double-stranded RNA can be processed into ∼21–23 nt small RNAs that direct sequence-specific cleavage of target transcripts, establishing the mechanistic basis of RNA interference [3, 4]. miRNAs provide an additional layer of post-transcriptional regulation and can influence virus-host interactions, including in insects [5, 6]. Small-RNA pathways also contribute to genome defence through transposon control [7, 8] and antiviral defence across diverse eukaryotes [9, 10].

In insects, the siRNA pathway is a major antiviral defence [11-14]. During infection, viral double-stranded RNA is processed by Dicer-2 into virus-derived small interfering RNAs (vsiRNAs) [15-17]. These vsiRNAs are loaded into Argonaute 2, which directs sequence-specific cleavage of viral RNA [18, 19].

PIWI-associated pathways are best known for transposon control [7], with somatic transposon-targeting piRNAs widespread across arthropods [8]. In some arthropods, these pathways also generate virus-derived piRNAs or piRNA-like small RNAs [20-24]. Endogenous viral elements (EVEs), viral sequences integrated into host genomes, provide an additional source of virus-related small RNAs. EVEs can occur within piRNA-producing genomic regions and generate piRNAs [25, 26]. Experimental studies show that EVE-derived piRNAs can contribute to sequence-specific restriction of cognate viral RNA [27, 28]. Genome-encoded piRNAs can also initiate responder and trailer piRNA production following cleavage of complementary viral RNA [29].

These pathways produce characteristic signatures in small-RNA sequencing data, although virus-derived profiles vary among host-virus systems [10, 30]. Dicer-2-derived vsiRNAs typically form discrete ∼21-nt populations [15-17] and can display complementary duplex-overlap patterns consistent with Dicer cleavage [16, 17]. By contrast, piRNAs are generally longer (∼24–30 nt), often show 1U enrichment, and during ping-pong amplification form complementary pairs with a characteristic 10-nt 5′ overlap and 10A enrichment in the partner population [7, 31]. Similar signatures occur in virus-derived piRNA populations in mosquitoes [20, 22-24].

Extracting these signatures from sequencing data requires reproducible read mapping and analysis across read-length classes, strands and biological replicates. Many established workflows still rely on command-line alignment followed by separate downstream analysis in R, Bioconductor or other software environments. These workflows generate multiple intermediate files and can be difficult to use without bioinformatics experience.

viRome provides specialised analysis of viral sRNAs but requires pre-mapped BAM files as input [32]. The sRNAtoolbox web server, including sRNAbench, provides browser-based small RNA profiling and differential expression analysis [33], but browser-based use requires sequencing data to be uploaded for server-side processing. Local deployment is available but requires a separate software environment. MISIS-2 provides a graphical interface for viewing strand- and size-resolved sRNA maps along a viral reference, and additionally supports consensus sequence reconstruction and single-nucleotide polymorphism detection [34], but is a standalone Java application operating on pre-aligned SAM/BAM files.

To address these limitations, we developed sRNAMap, a single-file HTML5/JavaScript application that combines read mapping with small-RNA-specific analysis entirely within the browser. It accepts raw FASTA/FASTQ reads or pre-aligned BAM files and provides strand-resolved coverage and read-end profiles, read-length and nucleotide-bias profiling, and analyses of read-overlap signatures, phasing, non-templated additions and sequence diversity metrics. No installation or external runtime environment is required, and all computation occurs locally without server-side data upload. Analyses can be saved and restored as JSON or exported as self-contained HTML reports containing figures and analysis metadata.

## Methods

### Small RNA sequencing datasets and reference sequences

Benchmarking used two small RNA-seq replicate libraries from Semliki Forest virus (SFV4)-infected *Aedes aegypti* Aag2-AF319 Dcr2-knockout cells transiently transfected with wild-type Dcr2, previously generated by Gestuveo et al. [16] (SRA accessions SRR13810521 and SRR13810522). Raw sequencing reads were downloaded from the NCBI Sequence Read Archive and trimmed as described previously [16]. The same pre-processed reads were used for all sRNAMap and Bowtie 2 benchmarking. Reads were mapped to the SFV4 genome (GenBank accession KP699763; 11,445 nt). The two uncompressed replicate FASTQ files were 879 MB and 805 MB and contained 10,201,151 and 9,368,007 reads, respectively (19,569,158 reads in total).

### sRNAMap mapping algorithm

sRNAMap maps sRNA reads to reference FASTA sequences using a frequency-weighted k-mer seeding strategy. Each read and its reverse complement are queried against the reference. K-mers sampled at intervals of ⌊k/2⌋ nucleotides contribute inverse-frequency-weighted support to candidate loci, thereby down-weighting repetitive seeds. Candidate loci meeting a user-defined minimum seed-support threshold are ranked by cumulative support, and up to the 10 highest-ranking candidates are evaluated by ungapped end-to-end alignment. Alignment scores are calculated as:

read length − (mismatch count × mismatch penalty)

Alignments exceeding the user-defined mismatch allowance are discarded, with an optional filter to exclude alignments containing a 3′-terminal mismatch. If primary seeding identifies no candidate loci and adaptive seeding is enabled, the read is re-queried using a reduced k-mer size (k − 2; minimum 8 nt) sampled at every position. Users can configure k-mer size, minimum seed support, mismatch allowance and penalty, read-length filtering and multi-mapping behaviour.

Multi-mapping reads can be handled using one of three policies. The default “best” policy reports a single top-scoring alignment while recording the number of equally best alignments among the evaluated candidates. “All equally best” reports tied best alignments up to a user-defined maximum-loci setting, whereas “random best” reports one randomly selected alignment from the same capped set. The default maximum-loci setting is 10. All benchmarking used the default “best” policy.

Claude Opus 5 (Anthropic; used during February–March 2026) was used to assist with code development and debugging from an initial HTML implementation written by R.H.P.

### Browser performance benchmarking

Runtime benchmarking was performed on a Windows 10 Education workstation (x64) equipped with an Intel Xeon E5-1607 v4 CPU at 3.10 GHz and 16 GB RAM. sRNAMap was tested in Microsoft Edge (v143.0.3650.80, 64-bit) and Google Chrome (v143.0.7499.110, 64-bit) using four Web Worker threads.

Benchmarking used the default mapping parameters: a 15-nt k-mer size, minimum seed-support threshold of 1, ≤2 mismatches, mismatch penalty of 2, adaptive seeding enabled, an 18–30 nt read-length range and the “best” multi-mapping policy. Adapter clipping and quality filtering within sRNAMap were disabled because the input libraries had already been pre-processed as described above. Each browser and compression condition was benchmarked across 10 independent runs. Uncompressed and gzip-compressed versions of the same FASTQ files were tested separately.

### Bowtie 2 comparison

To assess whether sRNAMap recapitulates conventional short-read alignments, we benchmarked mapping output against Bowtie 2 [35] using parameters chosen to approximate sRNAMap’s default mapping parameters. Prefiltered 18–30 nt reads were mapped using Bowtie 2 (v2.4.5) in end-to-end mode with an exact seed length of 15 nt `(-L 15, -N 0),` approximating sRNAMap’s default 15-nt primary seeding. To approximate sRNAMap’s default mismatch tolerance (≤2 mismatches) and mismatch penalty, Bowtie 2 was configured with mismatch penalties (`--mp 2,2`) and a minimum score threshold `(--score-min L,-4,0).` Bowtie 2 was permitted to report up to 10 loci per read `(-k 10),` with only primary alignments used for downstream comparison.

Bowtie 2 alignments were converted to sorted BAM files and filtered using SAMtools (v1.16.1) [36]. Strand-aware per-base coverage and 5′- and 3′-endpoint counts were calculated from primary alignments to generate coordinate-matched profiles for total read depth and strand-resolved read ends. The two replicate libraries were combined for this comparison. Concordance between sRNAMap and Bowtie 2 was evaluated at each SFV4 genome position using linear regression in GraphPad Prism v10.4.1.

## Results

### Tool implementation and architecture

sRNAMap is implemented as a single-file HTML5/JavaScript application (<1 MB) that runs entirely in the browser (Figure 1). Reads are loaded locally from FASTA/FASTQ files (uncompressed or gzip-compressed), and mapped against reference FASTA sequences. For FASTQ input, users can optionally perform 3′ adapter clipping and Phred-based quality filtering before mapping, with configurable adapter overlap, quality encoding, minimum quality and post-trimming read length. Alternatively, BAM alignments can be imported directly for downstream analysis and visualisation without remapping. Multiple BAM files can be analysed as replicate libraries, with configurable read-length, mapping-quality, primary-alignment and paired-read filters. sRNAMap parses BAM files directly in the browser, allowing externally aligned libraries to enter the same downstream analysis and visualisation. sRNAMap uses Web Workers to parallelise raw-read mapping and reduce blocking of the main user interface. For BAM workflows, a reference FASTA is optional but enables reference-sequence display, nucleotide-context analyses and reference-based validation of terminal soft-clips during NTA analysis. Multiple FASTQ or BAM files can be analysed as replicate libraries, with replicate-aware summaries available throughout the downstream analysis modules.

**Figure 1:**
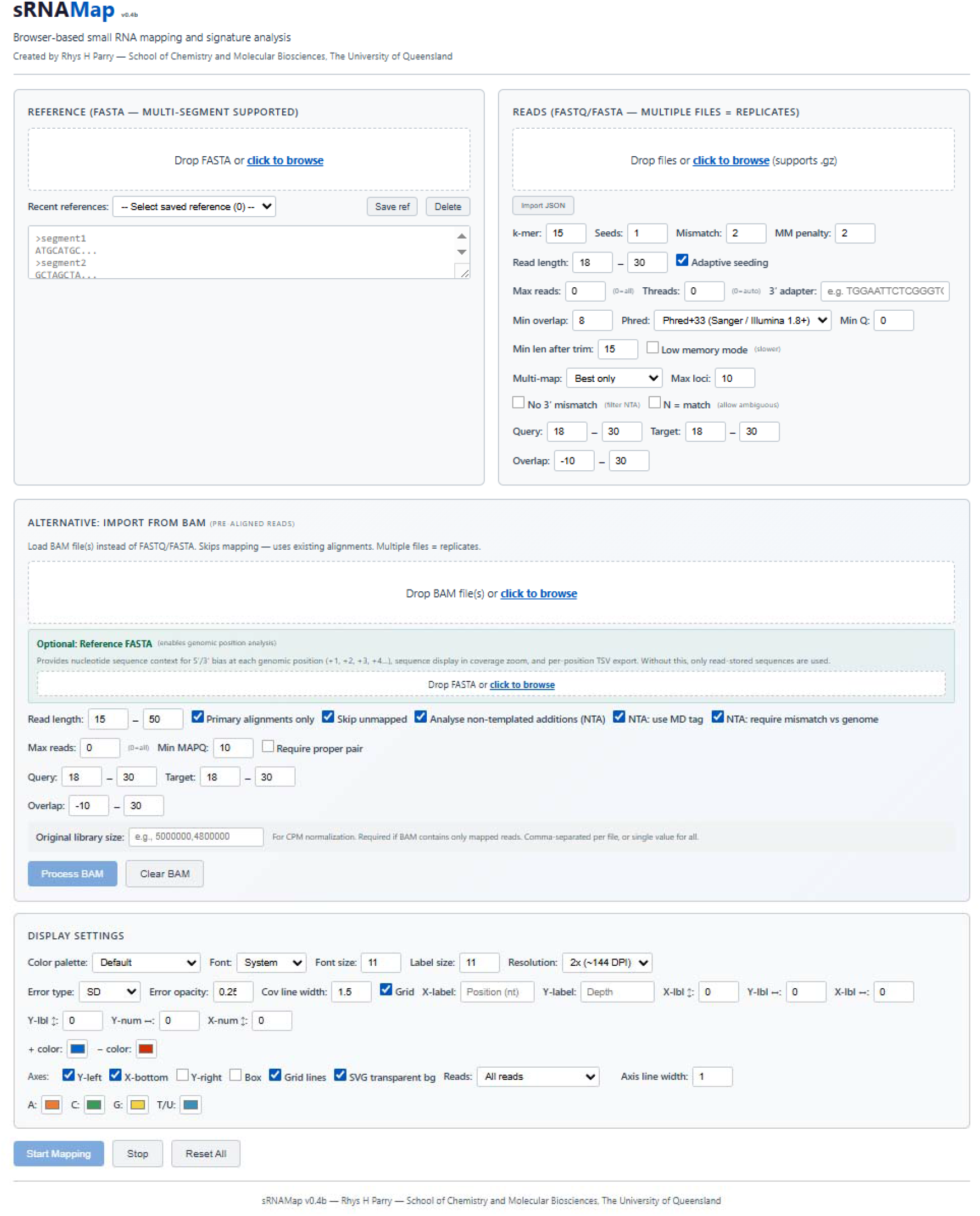
The sRNAMap user interface. Screenshot of the sRNAMap HTML web application showing the two primary input workflows. Users can map FASTA/FASTQ reads directly against reference FASTA sequences or import one or more pre-aligned BAM files for downstream analysis without remapping. Optional preprocessing, mapping, BAM filtering and display settings can be configured before analysis. Results can be exported and re-imported as JSON.

To support reproducible re-analysis, sRNAMap can export analysis results and settings as a JSON file, including length distributions, coverage arrays, overlap/signature matrices, and key parameter settings. These files can later be re-imported to regenerate plots and tables without repeating the analysis.

### Performance benchmarking

Across 10 repeated runs, the mean wall-clock runtime for the two SFV4 libraries was 139.17 s (2 min 19 s) in Microsoft Edge, corresponding to approximately 140,600 reads processed per second. A total of 1,788,852 reads mapped to SFV4 (9.1%), equivalent to approximately 12,850 mapped reads per second. Chrome was slower, with a mean runtime of 224.37 s (3 min 44 s), although mapped read counts were identical between browsers. Gzip compression had little effect on runtime, with mean mapping times of 139 s in Edge and 222 s in Chrome.

## Analysis modules

### Library-wide read-length and terminal nucleotide profiles

sRNAMap summarises read-length distributions and terminal nucleotide usage across user-defined size ranges (Figure 2A). These profiles help distinguish sRNA populations, including discrete 21–22 nt siRNA-like peaks and broader 24–30 nt piRNA-like distributions. For replicate libraries, the module reports mean distributions with user-selected error estimates (SD, SEM or 95% CI) and library size normalisation as counts per million (CPM).

**Figure 2:**
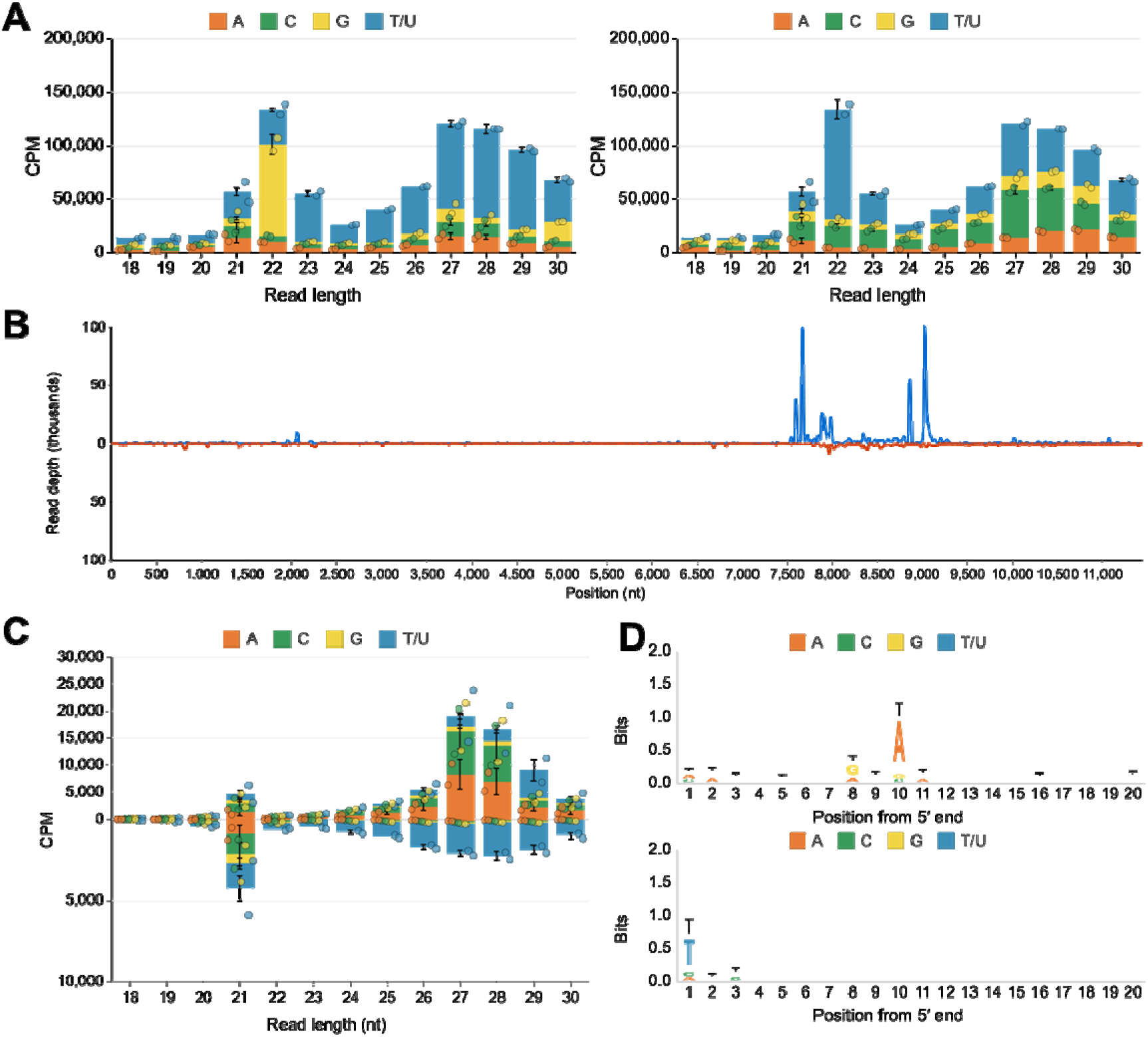
sRNAMap visualisations of the SFV4 benchmarking dataset. Small RNA libraries were derived from SFV4-infected *Ae. aegypti* Aag2-AF319 Dcr2-knockout cells transiently transfected with wild-type Dcr2 [16] and mapped to the SFV4 genome (GenBank accession KP699763). **(A)** Read-length distributions stratified by 5′ (left) and 3′ (right) terminal nucleotide identity, shown as counts per million. **(B)** Strand-resolved coverage of 18–30-nt reads across the SFV4 genome, with sense (+) coverage above and antisense (−) coverage below the x-axis. **(C)** Read-length distribution of SFV4-mapping reads stratified by 5′ nucleotide identity, showing U enrichment among antisense-mapping 24–30-nt reads. **(D)** Positional nucleotide-bias analysis of 24–30-nt reads. Sense- and antisense-mapping reads are shown in the upper and lower panels, respectively.

### Coverage profiling and visualisation

sRNAMap generates genome-wide coverage plots with sense (+) and antisense (−) reads displayed on opposite axes to show strand bias and coverage hotspots (Figure 2B). The supplied FASTA sequence defines the sense reference orientation; reads mapping to the opposite strand are reported as antisense (−), so a separate reverse-complement reference is not required. In addition to per-base coverage, sRNAMap calculates and exports the 5′ (start) and 3′ (end) positions of mapped reads, which can be used to examine Dicer cleavage patterns, piRNA phasing and processing boundaries. The coverage viewer supports interactive zoom and panning, and can reveal the underlying reference sequence at high zoom for examining local motifs or boundaries that coincide with sRNA coverage peaks.

### Mapped read length histogram and nucleotide bias analysis

sRNAMap displays mapped read-length distributions according to 5′- or 3′-terminal nucleotide identity (Figure 2C). This can distinguish a 21-nt siRNA-like population with minimal terminal bias from a 27–29-nt piRNA-like population with strong 1U enrichment, and allows comparison across references, segments or experimental conditions. Users can additionally generate positional nucleotide-bias profiles for selected read-length and strand classes (Figure 2D). In the SFV4 dataset, 24–30-nt sense-mapping reads show A enrichment at position 10, whereas antisense-mapping reads show strong 1U enrichment.

### Read-overlap signatures and phasing analysis

Different small RNA biogenesis pathways produce characteristic read-overlap signatures. In *Ae. aegypti*, 21-nt vsiRNAs can form duplexes with 19-nt complementarity and 2-nt 3′ overhangs. Intact vsiRNA duplexes accumulate when Ago2 passenger-strand slicing is impaired, increasing the 19-nt overlap signal [17]. In contrast, piRNAs generated through ping-pong amplification display a distinctive 10-nt 5′-to-5′ overlap between sense and antisense reads [20, 23].

sRNAMap quantifies these signatures using four end-comparison modes: 5′(+) vs 5′(−) for ping-pong detection, 3′(+) vs 3′(−) for 3′ end coordination, and 5′(+) vs 3′(−) or 3′(+) vs 5′(−) for cross-strand end relationships. The module reports raw or library-normalised overlapping read-pair counts, enrichment z-scores, overlap probabilities and probability-based z-scores (Figure 3A). The probability and probability z-score calculations reimplement the h-signature approach of Antoniewski [37] in JavaScript, following the reference signature.py implementation. A read-length-by-overlap heatmap shows whether overlap signatures are associated with particular read-length classes, such as ping-pong peaks restricted to piRNA-length reads versus broader patterns indicative of other processing modes (Figure 3B). The module can also express the same read pairs as stepRNA-style overhang and underhang distances [38]. In this representation, zero indicates a blunt end, negative values indicate an overhang and positive values an underhang. A canonical 2-nt Dicer signature appears directly as a −2 peak.

**Figure 3:**
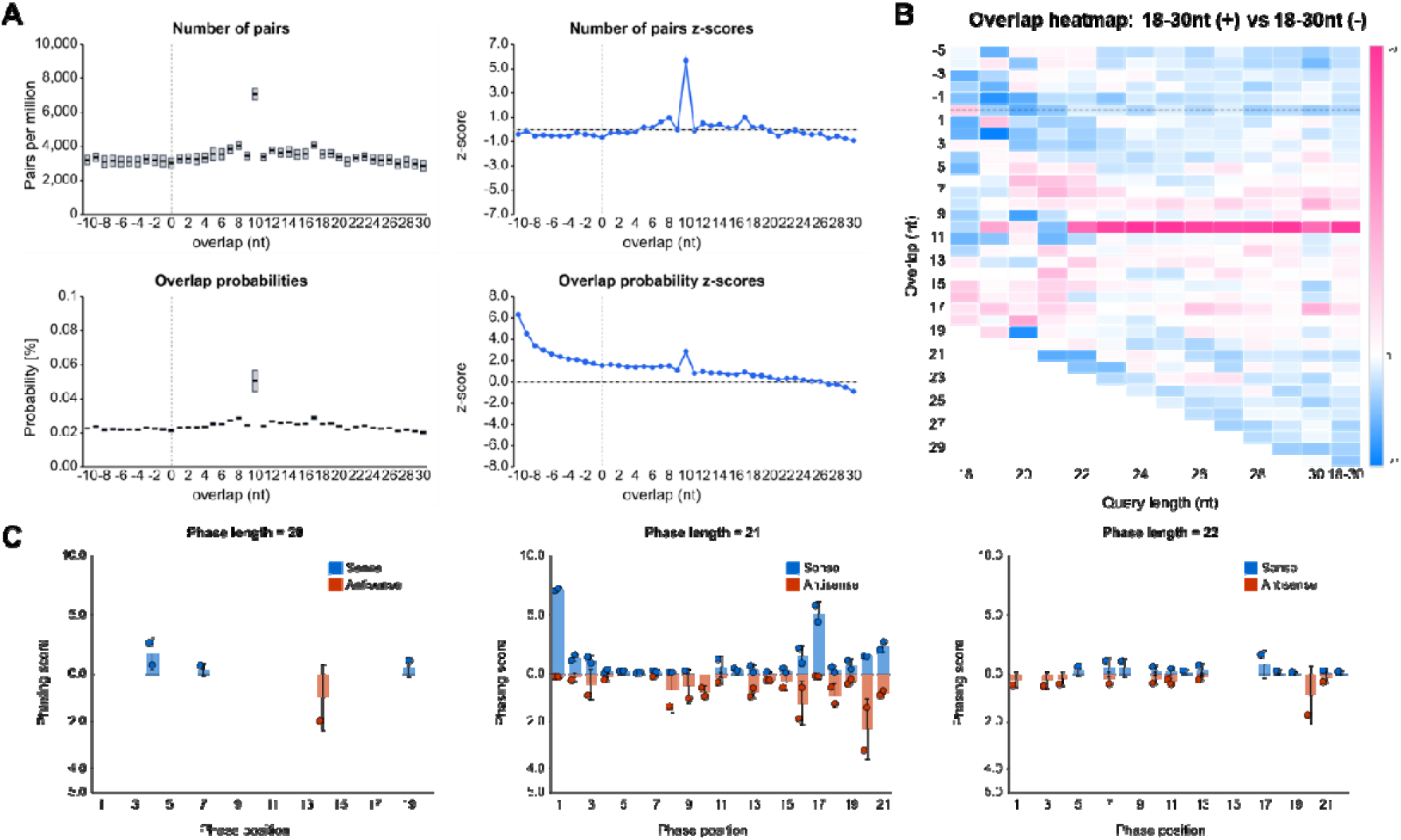
Overlap signature and phasing analysis outputs for the SFV4 benchmarking dataset. **(A)** Four-panel summary of 5′-to-5′ overlaps between sense (+) and antisense (−) mapped reads (18–30 nt). Top left: overlapping read pairs normalised to pairs per million; top right: enrichment z-scores; bottom left: overlap probabilities; bottom right: probability-based z-scores (h-signature). Data are shown as range charts with the mean indicated. **(B)** Read-length-by-overlap heatmap showing overlap signatures across sRNA size classes. The 10-nt ping-pong signature is strongest among 27–29-nt reads; the colour scale represents probability z-scores. **(C)** Phasing analysis of 20-, 21- and 22-nt reads, with phase length matched to read length. Phasing scores were calculated for each replicate and are shown as means.

sRNAMap also includes a phasing analysis module adapted from the phasing score described by Remnant et al. [39] to detect periodicity in small RNA production. In plants, phased siRNA biogenesis can be initiated by miRNA-guided cleavage that sets the register for subsequent Dicer-like processing [40]. In our SFV4 benchmarking dataset, no clear dominant phasing signal was observed across 20-, 21- or 22-nt reads mapping to the viral genome (Figure 3C). The module allows users to test user-defined read lengths and phase intervals for periodic processing signatures, and to restrict the analysis to a defined genomic interval. Because a phasing signal is only detectable when the analysis window is in the correct register relative to the initiating cleavage site, the module includes an offset scan that evaluates every phase register for the selected phase length and reports the register giving the strongest signal, which can then be applied to the main analysis. A flat offset-scan profile indicates that no single-phase register is dominant. A read-length-by-phase-position heatmap shows phasing and anti-phasing patterns across sRNA size classes.

### Non-templated addition (NTA) analysis

Small RNAs can carry non-templated nucleotides at their 3′ and, less commonly, 5′ termini, with 3′ uridylation in particular associated with altered stability, Argonaute loading and target repression [41]. For BAM input, sRNAMap identifies candidate non-templated additions from terminal soft clips and, where available, terminal mismatches encoded in the MD tag. When a reference FASTA is available, soft-clipped sequence that fully matches the corresponding reference sequence is excluded as templated, reducing false-positive NTA calls. The module reports the proportion of tailed reads, addition length and nucleotide composition, read-length distributions and motif-by-length summaries, with replicate-aware CPM normalisation and SVG, PNG and TSV export. Reads can be grouped as NTA-negative, NTA-positive or all reads, allowing comparison of coverage, read length, nucleotide-bias and other downstream profiles between non-tailed and tailed populations.

### Sequence diversity analysis

sRNAMap includes a sequence-diversity module for quantifying diversity among mapped sRNA sequences. For each selected read population, the module calculates Shannon diversity (H′ = −Σp□lnp□), Simpson diversity (1 − Σp□^2^), sequence richness (S; the number of unique sequences) and evenness (H′/lnS). Metrics can be stratified by replicate, read length or strand. Because sequence-level diversity requires enumeration of unique reads, this analysis can be memory intensive for large datasets. Sequence richness is also dependent on sampling depth and should therefore be compared between libraries in the context of library size.

### Flanking nucleotide bias

Previous work has identified characteristic nucleotide biases immediately flanking piRNA termini, including enrichment for U at the +1-position associated with phased piRNA biogenesis [29]. sRNAMap examines reference-sequence composition immediately flanking mapped sRNA termini. The module calculates nucleotide frequencies at positions −2 and −1 upstream and +1 and +2 downstream of mapped reads and standardises these values across read-length classes using z-scores, separately by strand. This enables flanking sequence-context signatures to be compared across sRNA size classes.

### Benchmarking against Bowtie 2

Coverage profiles across the SFV4 reference showed highly similar patterns between sRNAMap and Bowtie 2 (Figure 4A). Per-position values were highly concordant for total read depth (R^2^ = 0.9983), 5′-end counts (R^2^ = 0.9720) and 3′-end counts (R^2^ = 0.9688) (Figure 4B).

**Figure 4:**
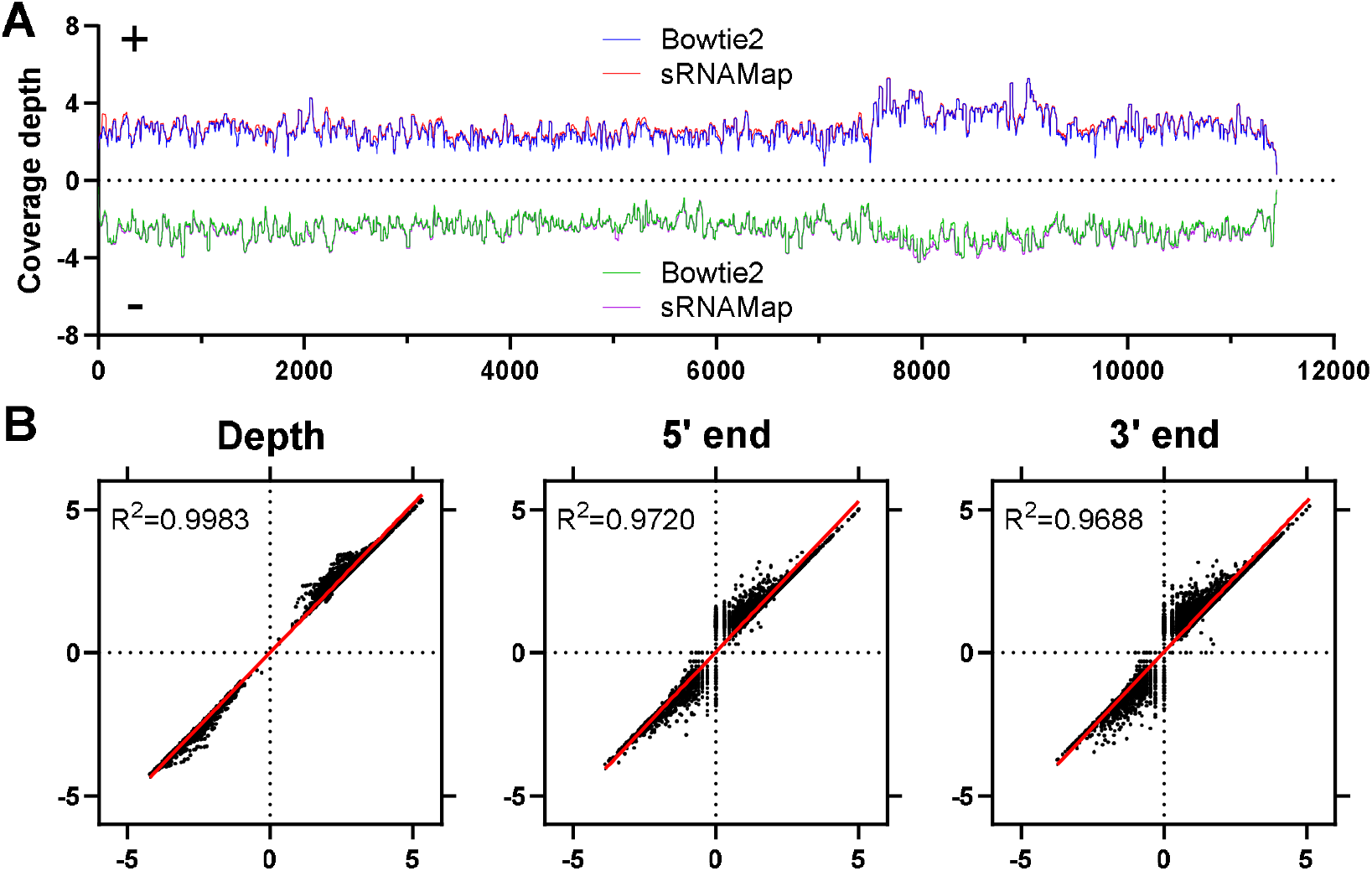
Concordance of strand-resolved coverage profiles between sRNAMap and Bowtie 2. **(A)** Genome-wide coverage showing strand-separated depth across the SFV4 genome as described above (positive axis: sense-strand coverage; negative axis: antisense-strand coverage). Bowtie 2 profiles are shown in blue/green and sRNAMap profiles in red/magenta, with the dashed line indicating zero coverage. **(B)** Per-position concordance between sRNAMap and Bowtie 2 for strand-resolved total depth, 5′ endpoint counts and 3′ endpoint counts, with sense-strand values plotted as positive and antisense-strand values as negative. Each point represents a single SFV4 genome position. The red line denotes a linear regression fit, and the coefficient of determination (R^2^) is shown in each panel. Reads were restricted to 18–30 nt for both tools, and Bowtie 2 was configured to approximate the sRNAMap default settings. The two replicate libraries were combined for this comparison.

### Limitations

Because sRNAMap performs all processing in the browser, available CPU and memory can limit analysis of very large datasets. sRNAMap provides optional in-browser 3′ adapter trimming and Phred-based quality filtering; however, pre-processed FASTQ files are recommended for very large libraries to reduce runtime. Base-quality scores can be used during preprocessing but are not incorporated into alignment scoring. sRNAMap encodes mapping coordinates in a packed representation supporting a maximum of 67 Mb (2^2^[bp) per reference sequence or segment. This accommodates viral genomes, transcript sets, transposable element consensus sequences and many genomic scaffolds, but individual assembled chromosomes exceeding this length must be split before analysis. Sequence-diversity analysis can be particularly memory intensive because it requires enumeration of unique read sequences. Because the built-in mapper uses ungapped end-to-end alignment, reads with substantial terminal additions may fail to map. For NTA-focused analyses, BAM files generated using local alignment, for example from Bowtie 2 in `--local` mode, are recommended. Performance and mapping concordance were evaluated using two replicate sRNA libraries and a single viral reference genome; performance will vary with library size, reference complexity and available hardware.

## Conclusions

sRNAMap combines direct FASTA/FASTQ mapping and analysis of pre-aligned BAM files in a single browser application. It provides strand-resolved coverage, read-length and nucleotide-bias profiling together with analyses of overlap signatures, phasing, terminal additions, sequence diversity and flanking nucleotide context. Benchmarking against Bowtie 2 demonstrated highly concordant per-position depth and read-end profiles, while direct FASTA/FASTQ mapping avoids the need to generate intermediate SAM/BAM files for browser-based analysis. JSON export enables analyses to be saved, re-imported and shared. sRNAMap is reference-agnostic and can be applied to viral genomes, cellular transcripts, transposable elements, synthetic constructs and other user-defined references within the per-sequence length limit described above.

### Software availability and requirements

sRNAMap is distributed as a single self-contained HTML file and is available at https://github.com/rhparry/sRNAMap under the MIT licence. The software version described and benchmarked in this study is sRNAMap v0.4b. sRNAMap requires no installation, runtime environment or server-side component and runs in modern browsers supporting Web Workers. It was tested in Microsoft Edge and Google Chrome (see Methods). All computation is performed locally and sequencing data are not transmitted from the user’s machine. sRNAMap uses the browser-native DecompressionStream API for gzip decompression where available and bundles fflate v0.8.2 (https://github.com/101arrowz/fflate; MIT licence) as a fallback. No other third-party dependencies are used.

## Supporting information

sRNAMap Stable Release

## Data summary

sRNAMap v0.4b is available under the MIT licence at https://github.com/rhparry/sRNAMap. The small RNA-seq datasets used for benchmarking are available from the NCBI Sequence Read Archive under accessions SRR13810521 and SRR13810522, with the SFV4 reference genome available under GenBank accession KP699763. Supporting benchmarking files are available from Figshare at https://doi.org/10.6084/m9.figshare.30951428. No new sequencing data were generated in this study.

## Author contributions

Conceptualisation, R.H.P.; Investigation, M.R., L.D.H., C.C.W.A., W.W., and R.H.P.; Visualisation, R.H.P.; Formal analysis, R.H.P.; Software, R.H.P.; Resources, E.S., A.K., R.H.P.; Validation, M.R., L.D.H., C.C.W.A., G.A.K., W.W., E.S., A.K., R.H.P.; Writing – original draft, R.H.P.; Writing – review and editing, M.R., L.D.H., C.C.W.A., W.W., G.A.K., E.S., A.K., R.H.P.

## Acknowledgements

R.H.P. is supported by NHMRC Ideas Grants (2038097 and 2049141) and by an Australian Research Council Discovery Early Career Researcher Award (DE260101042). This work was supported by resources provided by The University of Queensland Research Computing Centre’s Bunya supercomputer (doi:10.48610/wf6c-qy55), with funding from The University of Queensland, Brisbane, Australia.

