## Supplementary material for "sRNAMap: a lightweight, browser-based web application for small RNA mapping, analysis and visualisation": sRNAMap Stable Release: sRNAMap_v0.4b_stable_20_8_26 - Copy.html


### sRNAMap v0.4b

##### Reference (FASTA — multi-segment supported)

Drop FASTA or **click to browse**

Recent references:

-- Select saved reference --

Save ref
Delete

##### Reads (FASTQ/FASTA — multiple files = replicates)

Drop files or **click to browse** (supports .gz)

Import JSON

k-mer:

Seeds:

Mismatch:

MM penalty:

Read length:–

Adaptive seeding

Max reads:(0=all)

Threads:(0=auto)

3′ adapter:

Min overlap:

Phred:Phred+33 (Sanger / Illumina 1.8+)Phred+64 (Illumina 1.3-1.5)

Min Q:

Min len after trim:

Low memory mode(slower)

Multi-map:
Best only
All equally best
Random best

Max loci:

No 3′ mismatch(filter NTA)

N = match(allow ambiguous)

Query:–

Target:–

Overlap:–

##### Alternative: Import from BAM (pre-aligned reads)

Load BAM file(s) instead of FASTQ/FASTA. Skips mapping — uses existing alignments. Multiple files = replicates.

Drop BAM file(s) or **click to browse**

**Optional: Reference FASTA**
(enables genomic position analysis)

Provides nucleotide sequence context for 5′/3′ bias at each genomic position (+1, +2, +3, +4…), sequence display in coverage zoom, and per-position TSV export. Without this, only read-stored sequences are used.

Drop FASTA or **click to browse**

Read length:–

Primary alignments only

Skip unmapped

Analyse non-templated additions (NTA)

NTA: use MD tag

NTA: require mismatch vs genome

Max reads:(0=all)

Min MAPQ:

Require proper pair

Query:–

Target:–

Overlap:–

Original library size:

For CPM normalization. Required if BAM contains only mapped reads. Comma-separated per file, or single value for all.

Process BAM
Clear BAM

File 1/1
0 reads processed

Decompressing...
—

##### Display Settings

Color palette:
Default
Rhys
Pastel
Viridis
Colorblind-friendly

Font:
System
Arial
Helvetica
Times
Georgia
Courier

Font size:

Label size:

Resolution:
1x (~72 DPI)
2x (~144 DPI)
3x (~216 DPI)
4x (~288 DPI)
5x (~360 DPI)
8x (~600 DPI)

Error type:SDSEM95% CI

Error opacity:

Cov line width:

Grid

X-label:

Y-label:

X-lbl ↕:

Y-lbl ↔:

X-lbl ↔:

Y-lbl ↕:

Y-num ↔:

X-num ↕:

+ color:

− color:

Axes:

Y-left

X-bottom

Y-right

Box

Grid lines

SVG transparent bg

Reads:All readsPerfect only (no NTA)NTA only

Axis line width:

A:

C:

G:

T/U:

Start Mapping
Stop
Reset All

Processing...

Total Reads

0

Mapped

0

Mapped %

0%

Avg. Depth

0×

Breadth

0%

Segments

0

Replicates

0

###### Per-replicate Mapping Summary

Download TSV

Shows mapped read counts for each input file (replicate) across all reference segments. Rows highlighted in red have no mapped reads.

##### 5′ Bias Length Histogram

⟲All readsPerfect onlyNTA onlyUnmapped onlySVGPNGTSV

W:

H:

Y-max:

Y-tick:

X-start:

X-tick:

Cap:

Length:–

Bar %:

Data:CountsCPMPercentage

Font:

Line width:

Grid

X-label:

Y-label:

X-lbl ↕:

Y-lbl ↔:

X-lbl ↔:

Y-lbl ↕:

Y-num ↔:

X-num ↕:

X-off:

Y-off:

Error

Points

Jitter

Pt opacity:

Legend

Leg font:

Leg pos:CenterLeftRight

Bold

Hover for details

##### 3′ Bias Length Histogram

⟲All readsPerfect onlyNTA onlyUnmapped onlySVGPNGTSV

W:

H:

Y-max:

Y-tick:

X-start:

X-tick:

Cap:

Length:–

Bar %:

Data:CountsCPMPercentage

Font:

Line width:

Grid

X-label:

Y-label:

X-lbl ↕:

Y-lbl ↔:

X-lbl ↔:

Y-lbl ↕:

Y-num ↔:

X-num ↕:

X-off:

Y-off:

Error

Points

Jitter

Pt opacity:

Legend

Leg font:

Leg pos:CenterLeftRight

Bold

Hover for details

Distribution of all reads in library by length, colored by 5′ or 3′ terminal nucleotide. Useful for identifying siRNA (21nt, U-bias) vs piRNA (24-30nt, U-bias) populations. Shows total library composition regardless of mapping. **Note:** Both panels automatically use the same Y-axis scale for easy comparison.

**BAM import:** These histograms show mapped reads only (not full library). If your BAM contains only aligned reads, this represents mapped read distribution, not input library composition.

##### Coverage (+ sense up, − antisense down)

⟲All readsPerfect onlyNTA onlySVGPNGTSV

W:

H:

+Y max:

−Y max:

Segment:

Data:CountsCPM

Type:Full coverage5′ end only3′ end only

Font:

Gap:

Title:Chromosome nameCustom titleNo title

Min len:

Max len:

X-off:

Y-off:

X-tick:

X-start:

Y-tick:

Bin:

X-axis bold

Y-axis bold

Error

Show sequence

Lock Y-axis

Start:

End:

◀
−
+
▶
Reset

Drag to pan

Hover for position details

Genome-wide read coverage profile. Sense (+) reads shown above the axis, antisense (−) below. Hotspots indicate highly targeted regions.

##### Mapped Length Distribution

⟲All readsPerfect onlyNTA onlyUnmapped onlySVGPNGTSV

W:

H:

+Y max:

Fwd split %:

−Y max:

Y-tick:

Length:–

Segment:

Title:Chromosome nameCustomNo title

Top (fwd):5′3′

Bottom (rev):5′3′

Bar %:

Data:CountsCPMPercentage

X-tick:Every 1Every 2Every 5

Font:

Line width:

Grid

X-start:

X-label:

Y-label:

X-lbl ↕:

Y-lbl ↔:

X-lbl ↔:

Y-lbl ↕:

Y-num ↔:

X-num ↕:

X-off:

Y-off:

Error

Points

Jitter

Pt opacity:

Legend

Leg font:

Leg pos:CenterLeftRight

Bold

Cap:

Hover for count details

Strand-separated length distribution with nucleotide bias. Forward reads (top) and reverse reads (bottom) shown separately to reveal strand-specific patterns.

##### Mapped Nucleotide Bias

⟲All readsPerfect onlyNTA onlySVGPNG

W:

H:

Segment:

Title:NameCustomNone

End:5′3′

Strand:BothForward (+)Reverse (−)

Length:–

Display:ProportionBits (logo)

Max bits:

X-start:

X-tick:Every 1Every 2Every 5

Font:

Line width:

Grid

X-label:

Y-label:

X-lbl ↕:

Y-lbl ↔:

X-lbl ↔:

Y-lbl ↕:

Y-num ↔:

X-num ↕:

Error

Legend

Leg font:

Leg pos:CenterLeftRight

Bold

Leg X-off:

Leg Y-off:

Hover for nucleotide details

Positional nucleotide composition from 5′ or 3′ end of mapped reads. Strong 1U bias at position 1 indicates Argonaute loading. Position 10A bias suggests ping-pong amplification. Use length filter to focus on specific size classes (e.g., 21-21 for siRNA, 24-30 for piRNA).

##### Overlap Signatures (4-Panel Summary)

⟲Overlap (signature.py)stepRNA 3′ distancestepRNA 5′ distanceAll readsPerfect onlyNTA onlySVGPNGTSV

Uses logic from Antoniewski (2014) Methods Mol Biol, DOI: 10.1007/978-1-4939-0931-5\_12

W:

H:

Segment:

Font:

Line width:

Grid

X-label:

Y-label:

X-lbl ↕:

Y-lbl ↔:

X-lbl ↔:

Y-lbl ↕:

Y-num ↔:

X-num ↕:

End mode:
5′(+) vs 5′(−)
3′(+) vs 3′(−)
5′(+) vs 3′(−)
3′(+) vs 5′(−)

Query (+) min:

Query (+) max:

Target (−) min:

Target (−) max:

Overlap min:

Overlap max:

Recalculate

Error bars

Points

Normalize to RPM

Counts Y:

Prob Y:

z Y:

Counts Y tick:

Prob Y tick:

z Y tick:

X tick:

Panel width:

Panel height:

Title size:

Bold titles

Panel subtitles

X offset:

Y offset:

Panel borders

Show 10nt line

Hover for overlap details

Overlap analysis between sense (+) and antisense (−) reads using selected end positions. 5′(+) vs 5′(−) is the classic ping-pong signature (10nt peak). The asymmetric modes 5′(+) vs 3′(−) and 3′(+) vs 5′(−) explore cross-strand end relationships that may reveal novel processing patterns.

Antoniewski C. Computing siRNA and piRNA overlap signatures. *Methods Mol Biol.* 2014;1173:135-46. doi: 10.1007/978-1-4939-0931-5\_12. PMID: 24920366

##### Overlap Heatmap (by Query Length)

⟲TSVAll readsPerfect onlyNTA onlySVGPNG

Uses logic from Antoniewski (2014) Methods Mol Biol, DOI: 10.1007/978-1-4939-0931-5\_12

W:

H:

Font:

Segment:

Title:NameCustomNone

Cap overlap at shorter read

Data:Probability z-score (h-sig)Count z-scoreProbabilityCounts

Colors:Blue→PinkViridisPlasmaCool→Warm

X-tick:

Z min:

Z max:

Query (+) min:

Query (+) max:

Target (−) min:

Target (−) max:

Overlap min:

Overlap max:

Hover for heatmap details

Heatmap showing overlap signatures by query read length. Probability z-score (h-sig) is recommended as it normalizes per-position, making it robust to depth variations. Count z-score may be biased by high-coverage regions. Each row represents a different query length, with 'all' aggregating across lengths.

Antoniewski C. Computing siRNA and piRNA overlap signatures. *Methods Mol Biol.* 2014;1173:135-46. doi: 10.1007/978-1-4939-0931-5\_12. PMID: 24920366

##### Non-Templated Additions (NTA)

TSVBar SVGBar PNGHist SVGHist PNGHeat SVGHeat PNG

Untemplated 3'/5' tails from terminal soft-clips (local alignment) or MD-tag terminal mismatches (end-to-end). Requires BAM input.

End:3′ end5′ end

Segment:All segments

Bar W:

Bar H:

Hist W:

Hist H:

Font:

Line width:

Grid

Hist data:CPMCounts

Len min:

Len max:

X-tick:

X-start:

Y min:

Y max:

Y-tick:

Error bars

Bar %:

Motif heatmap:

Motif:Mono (4)Di (16)Tri (64)

Scale:CPM (mean per library)Raw counts% of motif (row)% within length (column)

Max rows:

Colour max:

Heat W:

Heat H:

Enable NTA analysis and re-import a BAM to populate.

Left: proportion of mapped reads carrying an untemplated tail, and the size class of that tail (mono / di / tri / tetra+). Right: length distribution of tailed reads, coloured by the identity of the terminal added nucleotide. Lengths are templated length (read length minus the untemplated tail).

##### Phasing Analysis

⟲All readsPerfect onlyNTA onlySVGPNG

Adapted from Remnant et al. (2017) J Virol, DOI: 10.1128/JVI.00158-17

W:

H:

Font:

Line width:

Grid

X-label:

Y-label:

X-lbl ↕:

Y-lbl ↔:

X-lbl ↔:

Y-lbl ↕:

Y-num ↔:

X-num ↕:

Segment:

Phase length:

Cycles:

Read min:

Read max:

Pos start:

Pos end:

Offset:

RPM normalize

Error bars

Points

+Y max:

−Y max:

Bar %:

Legend


Register Scan (find best offset)

**The Register Problem:** Phasing analysis assumes position 0 of your reference aligns with the start of a Dicer cleavage register.
However, for incomplete genomes or arbitrary reference coordinates, this alignment is unknown. A strong phasing signal at position 9
might simply mean your reference starts 9nt into the true biological register—not that position 9 is biologically special.

**Solution:** This scan iterates through all possible register offsets (0 to phase\_length−1), calculating the phasing score
at each offset. The offset producing the highest score indicates where Dicer processing likely begins relative to your reference.
For each offset *k*, all read positions are shifted by +*k* before calculating the standard phasing score at register position 1.

**Interpretation:** A clear peak at one offset suggests consistent processive Dicer activity starting at that register.
A flat profile suggests either no phasing or phasing that is not register-dependent. Apply the best offset to align your analysis
with the biological cleavage register.

Scan All Offsets

Apply Best
SVG
PNG

Click "Scan All Offsets" to find the best phase register


Hover for phasing details

Phasing analysis detects Dicer-mediated processing signatures. If small RNAs are produced by Dicer cutting at regular intervals along a dsRNA precursor, reads should accumulate at positions that are multiples of the phase length apart. High phasing scores at specific positions indicate structured, Dicer-dependent biogenesis. Use the **Register Scan** below to identify the optimal offset when your reference coordinates don't align with the biological cleavage start site.

**Score = ln[((1 + 10 × P) / (1 + U))^n−2^]** where P = in-phase reads (RPM-normalized if enabled), U = out-of-phase reads, n = number of occupied cycle positions. Positive scores indicate phasing; negative scores indicate anti-phasing.

##### Phasing Heatmap

⟲All readsPerfect onlyNTA onlySVGPNG

Heatmap showing phasing scores by read length and phase position. Positive scores (red) indicate evidence of phasing; negative scores (blue) indicate anti-phasing (more out-of-phase reads than expected). White indicates neutral/no signal. Phase length should typically be 21nt for plant siRNAs or the expected Dicer cleavage interval.

W:

H:

Font:

Segment:

Colors:
Blue-Red
Viridis
Plasma
Grayscale

Z min:

Z max:

Hover for details

##### Sequence Diversity

Calculate⟲All readsPerfect onlyNTA onlySVGPNGTSV

Shannon diversity (H') measures the complexity of the small RNA population. **Click "Calculate" to analyze** — this panel requires storing unique sequences which uses significant memory. Pre-calculated stats shown if available from streaming.

W:

H:

Font:

Segment:

Metric:
Shannon (H')
Simpson (1-D)
Richness (unique seqs)
Evenness (H'/ln(S))

Group by:
Replicate
Read length
Strand

Points

Error bars

Read min:

Read max:

Y min:

Y max:

Box %:

Click "Calculate" to compute diversity metrics

**H' = −Σ(p_i_ × ln(p_i_))** where p_i_ = proportion of reads from sequence i. Range: 0 (single sequence) to ln(S) where S = number of unique sequences.

##### Downstream Nucleotide Bias

⟲All readsPerfect onlyNTA onlySVGPNGTSV

Inspired by Joosten et al. (2021) Nucleic Acids Res, DOI: 10.1093/nar/gkab640

Z-score of reference nucleotide frequency at positions flanking mapped reads: −2, −1 (upstream of read start) and +1, +2 (downstream of read end) on the genome. Computed across read sizes within each strand. For sense piRNAs, the Zucchini +1U signature appears at +1; for antisense piRNAs it appears as +1A (= U on the antisense strand) at −1.

W:

H:

Font:

Segment:

Read min:

Read max:

Min reads/size:

Z cap:

Show values

Hover for nucleotide bias details

Each heatmap row = one read size. Columns are A, U, G, C grouped by genomic position (−2, −1 before read; +1, +2 after read). Red = enriched relative to other sizes; blue = depleted. For sense piRNAs (25–30 nt, bold), look for U enrichment at +1; for antisense piRNAs, look for A enrichment at −1 (complement of U on the antisense strand). The red line marks the read boundary. **Requires reference FASTA.**

Export Full Statistics (JSON)

##### Export Standalone Report

Generate a self-contained HTML file with all figures and analysis metadata. Ideal for supplementary materials in manuscripts - reviewers can open it in any browser without additional software.

Filename:

Export Standalone HTML

sRNAMap v0.4b — Rhys H Parry — School of Chemistry and Molecular Biosciences, The University of Queensland
